# LoopDetect: Comprehensive feedback loop detection in ordinary differential equation models

**DOI:** 10.1101/2020.11.15.383703

**Authors:** Katharina Baum, Jana Wolf

## Abstract

**Summary:** The dynamics of ordinary differential equation (ODE) models strongly depend on the model structure, in particular the existence of positive and negative feedback loops. LoopDetect offers user-friendly detection of all feedback loops in ODE models in three programming languages frequently used to solve and analyze them: MATLAB, Python, and R. The developed toolset accounts for user-defined model parametrizations and states of the modelled variables and supports feedback loop detection over ranges of values. It generates output in an easily adaptable format for further investigation.

**Availability and Implementation:** LoopDetect is implemented in R, Python 3 and MATLAB. It is freely available at https://cran.r-project.org/web/packages/LoopDetectR/, https://pypi.org/project/loopdetect/, https://de.mathworks.com/matlabcentral/fileexchange/81928-loopdetect/ (GPLv3 or BSD license).

## 1 Introduction

Ordinary differential equations (ODEs) are used to model various biological systems from signaling to metabolism. They describe the development of variables over time based on their dependencies and interactions. Feedback loops are circular regulations where a variable is regulating itself either directly (self-loop) or via interactions with other variables. Feedback loops are of high importance for the possible dynamics of ODE models: Negative feedback loops are required for oscillations and can cause robustness, positive feedback loops can amplify signals and can induce multiple stable steady states (Domijan and Pecou, 2012; Ferrell, 2013; Tyson, et al., 2003).

Already in small biological models it is difficult to detect all feedback loops by visual inspection of model sketches (Nguyen and Kholodenko, 2015), in detailed mechanistic biological models the number of feedback loops can be very high (Choi, et al., 2012). Consequently, computational methods are required for reliable, systematic analysis. Tools such as GinSim (Chaouiya, et al., 2012), CellNetAnalyzer (Klamt and von Kamp, 2009) or Cytoscape (Shannon, et al., 2003) that are dedicated to the analysis of logical models or graphs *per se* allow the determination of feedback loops. However, these are not capable of reporting the feedback loops in ODE models. There are standalone tools to determine the positive feedback loops in ODE models that cause bistability (Feliu and Wiuf, 2015) and/or other criteria than feedback loops for oscillations (Donnell, et al., 2014). These are based on chemical reaction network theory (CRNT) and thus target general properties of model structures. However, frequently, insights are desired at specific model parametrizations and solutions that fit to the biological observations described by the model.

LoopDetect is a tool suite complementary to CRNT-based approaches that provides comprehensive detection of feedback loops of ODE models at user-defined parametrizations and sets of variable values. It is implemented in three programming languages popular with ODE model analysis: MATLAB, Python, and R.

## 2 Results

Determining feedback loops with LoopDetect is based on the fact that the Jacobian matrix contains direct and directed interactions between variables and thus represents the ODE system’s underlying interaction graph (Thomas and Kaufman, 2002). For numerical Jacobian matrix determination, LoopDetect employs complex-step derivatives that decisively improve the exactness compared to normal finite differences (Martins, et al., 2003). This allows to distinguish zero entries of the Jacobian (non-existent regulations) from non-zero entries (present regulations). We leveraged established, efficient algorithms on circuit detection (Johnson, 1975) and strongly connected component detection (Tarjan, 1972) and adapted them to the situation of graphs generated from ODE models. Defaulting an upper limit to the number of returned feedback loops avoids memory exhaustion.

The output of LoopDetect is a list of loops containing: (i) the participating variables in the order given by the loop, (ii) the length of the loop, and (iii) its sign, positive or negative (see Fig. 1A-D for the example of a general biological oscillator model studied in (Baum, et al., 2016)). The list of loops is returned in established, popular data formats (MATLAB’s table, R’s data.frame, or Python’s pandas DataFrame, see Fig 1D). LoopDetect provides a function for efficient determination of feedback loops over sets of variable values (`find_loops_vset`). This feature determines changes in feedback loops over transient behavior or for different initial conditions. In addition, LoopDetect contains summary and comparison functions (Fig. 1E: comparison of two systems with either positive or negative regulation), and functions for querying loop lists for an edge of interest.

**Fig. 1.**
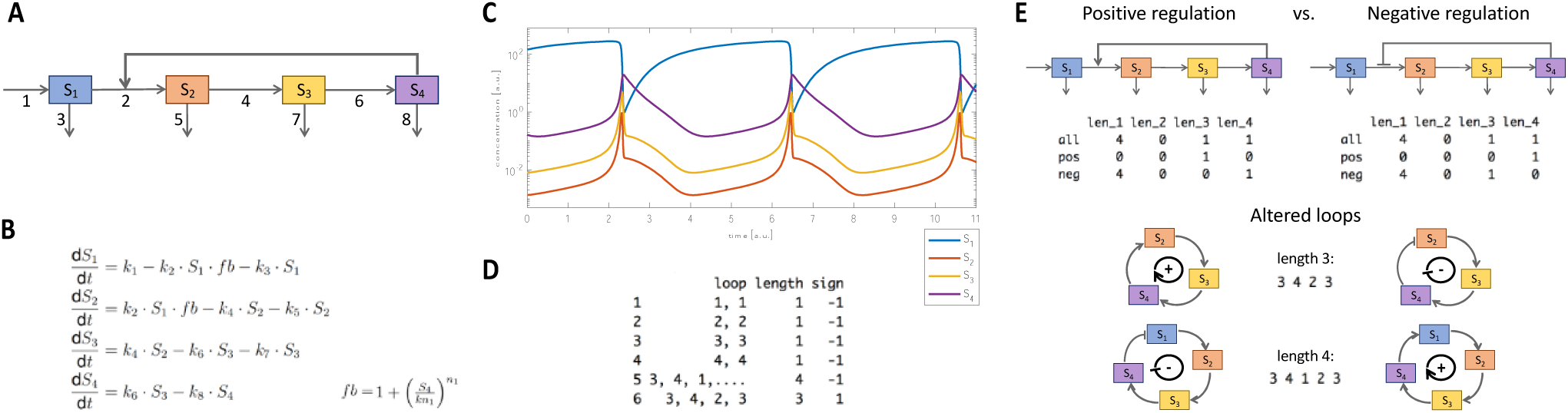
Example for feedback loop detection and comparison with LoopDetect. A-C: Sketch of a general oscillator model with positive feedback regulation (Baum, et al., 2016) (A), its corresponding ODE system (B) and the oscillatory solution at a specific parameter set (C). D: List of feedback loops (6 loops - 1 positive, 5 negative) detected with LoopDetectR in the model shown in (A), (B). E: Comparison of the feedback loops of the model with positive feedback regulation (left) to the same model where the feedback regulation is negative (right, 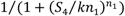), identical parameters). The summary tables of the feedback loops are generated by LoopDetectR. LoopDetect finds that four of the six feedback loops are identical in both systems while two differ (switched sign, visualized as altered loops).

We compared LoopDetect’s runtimes for processing Jacobian matrices between its three versions (Supplementary Material). The Python version detected feedback loops slightly faster than the MATLAB version. The R version of LoopDetect was faster than the other two for intermediate and larger systems (>30 variables) with many (ca. >100) feedback loops.

## 3 Conclusion

With its detailed documentation in MATLAB, Python, and R, LoopDetect provides a valuable, easy-to-use resource for comprehensive positive and negative feedback loop detection of ODE models based on graph theory algorithms. The tool suite enables detecting, analysis and comparison of feedback loops for model parametrizations and sets of variable values of interest to the user in the programming language they also use for parameter estimation, integration and plotting.

## Acknowledgements

We thank Sandra Krüger for valuable support in the initial stage of the project, and Oscar Arturo Migueles Lozano and Mareike Simon for testing the tools.

