## Supplementary Material for "LoopDetect: Comprehensive feedback loop detection in ordinary differential equation models"

### Runtime comparison of the R, MATLAB, Python versions of LoopDetect

We compared the runtimes of the LoopDetect versions in the three programming languages MATLAB, R, and Python. To this purpose, we sampled signed Jacobian matrices of different sizes (simulating systems with 30, 100, 300, 1000, or 3000 variables) with fixed numbers of non-zero interactions that enabled a broad range of numbers of feedback loops in the sampled systems. We measured the elapsed time for computing the list of feedback loops with each LoopDetect version as minimal value over three function calls for each input matrix. Thereby, we restricted the maximal number of detected and reported loops to $2.5x10^5$.
As is known for the employed algorithms from graph theory (Johnson's algorithm and Tarjan's algorithm), runtime increased with increasing number of variables (i.e. with system size) and with increasing number of feedback loops in the system for each of the three LoopDetect versions, LoopDetect_for_Matlab, LoopDetectR, and LoopDetect (Fig. S1A).
The Python version detected feedback loops slightly faster than the MATLAB version (Fig. S1B, C). While the R version of LoopDetect was faster than the other two for intermediate and larger systems with many feedback loops, both the MATLAB and the Python version outperformed the R version for small systems (30 variables) as well as for large systems (>300 variables) with few (up to ca. 100) feedback loops.

*More detailed specifications:*
We randomly sampled $10^3$, $10^3$, $10^3$, 500, or 250 matrices for system sizes of 30, 100, 300, $10^3$, or $3x10^3$ variables, respectively, with a fixed number of edges (non-zero entries of the matrix: 100, 200, 450, $1.3x10^3$, or $3.5x10^3$, respectively; values drawn from (-1,1)). Of those, we choose a subset to keep runtimes short: 200 matrices of size 30x30 (plus 50 with >$2.5x10^5$ loops), 400 matrices of size 100x100 (plus 50 with >$2.5x10^5$ loops), 913 matrices of size 300x300 (plus 50 with >$2.5x10^5$ loops), 419 matrices of size $10^3x10^3$ (plus 50 with >$2.5x10^5$ loops), 239 matrices of size 3000x3000 (plus 11 with >$2.5x10^5$ loops).
Computations were run on MacOS Mojave 10.14 on a 3.3 GHz Intel Core i7, 16 Gb 1867 MHz DDR3. We used MATLAB 2019b and the *tic* and *toc* commands for time measurements; R version 3.6.1 (2019-07-05), igraph_1.2.4.1, and the elapsed time from *system.time*; and Python 3.8 with networkx 2.4, numpy 1.18.5, pandas 1.0.4. and *default_timer* in the timeit module from timeit_plus 0.0.3 for measuring runtime.

Note that we only measured the time to derive the loop list from the Jacobian matrix, not the time required to compute or to read in the Jacobian, nor to write the resulting loop list to a file.

### Figure S1

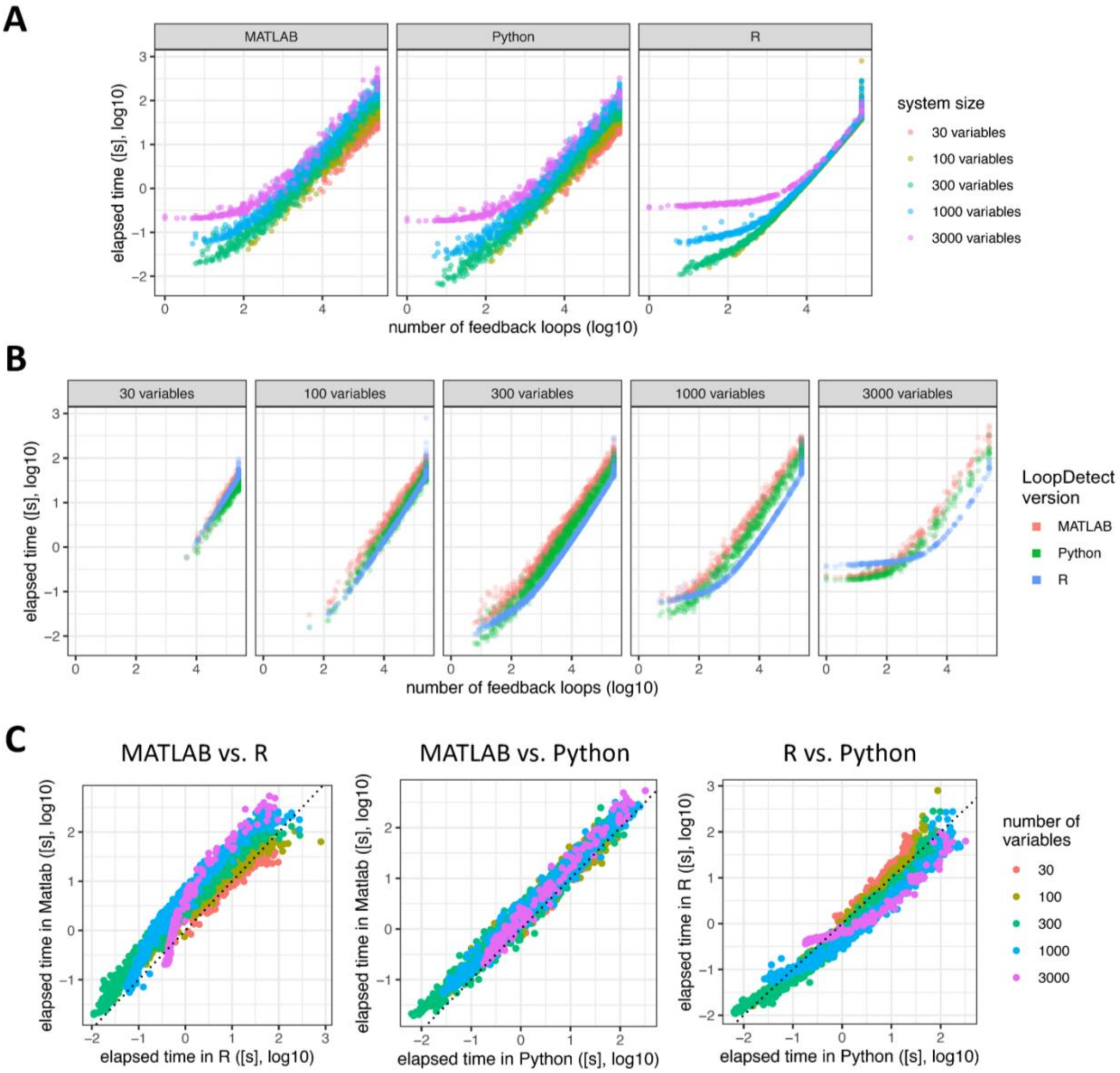


**A**: Runtimes for each LoopDetect version (left: MATLAB, center: Python, right: R) over the detected numbers of feedback loops. Each dot represents the runtime for one sampled Jacobian matrix. Colors denote the number of variables of the sampled system.
**B**: Runtimes of the LoopDetect versions (red: MATLAB, green: Python, blue: R) for different system sizes (increasing from left to right) depending on the number of detected feedback loops in the system.
**C:** Pairwise comparison of the runtimes of the LoopDetect versions (left: MATLAB vs. R, center: MATLAB vs. Python, right: R vs. Python) over all sampled matrices. Colors denote the system size.
